# In vivo proximity biotin ligation identifies the interactome of Egalitarian, a Dynein cargo adaptor

**DOI:** 10.1101/2021.07.01.450794

**Authors:** Frederick C. Baker, Hannah Neiswender, Rajalakshmi Veeranan-Karmegam, Graydon B. Gonsalvez

**Affiliations:** Cellular Biology and Anatomy, Medical College of Georgia, Augusta University, 1460 Laney Walker Blvd, Augusta, GA, 30912, USA

**Keywords:** RNA localization, molecular motor, cell polarity, nanobody, P body

## Abstract

Numerous motors of the Kinesin family contribute to plus-end directed microtubule transport. However, almost all transport towards the minus-end of microtubules involves Dynein. Understanding the mechanism by which Dynein transports this vast diversity of cargo is the focus of intense research. In select cases, adaptors that link a particular cargo with Dynein have been identified. However, the sheer diversity of cargo suggests that additional adaptors must exist. We used the *Drosophila* egg chamber as a model to address this issue. Within egg chambers, Egalitarian is required for linking mRNA with Dynein. However, in the absence of Egalitarian, Dynein transport into the oocyte is severely compromised. This suggests that additional cargoes might be liked to Dynein in an Egalitarian-dependent manner. We therefore used proximity biotin ligation to define the interactome of Egalitarian. This approach yielded several novel interacting partners, including P body components and proteins that associate with Dynein in mammalian cells. We also devised and validated a nanobody based proximity biotinylation strategy which can be used to define the interactome of any GFP-tagged protein.

## INTRODUCTION

Cellular polarity, a defining feature of eukaryotic cells, is achieved by the asymmetric sorting of mRNAs, proteins, vesicles, and organelles within the cell (Jossin, 2020). Often, these cargoes are sorted by transport on cytoskeleton filaments such as actin and microtubules. In general, actin-based transport is used for short-range movement and anchoring of cargo whereas the microtubule cytoskeleton and its associated motors are used for transporting cargo over longer distances.

Microtubules are inherently polarized and contain a minus-end that is usually anchored within a microtubule organizing center, and a plus-end that extends outward (Meiring et al., 2020). Most motors of the Kinesin family, of which there are forty five in humans, are involved in transporting cargo towards the plus-end of microtubules (Hirokawa et al., 2009). However, virtually all transport towards the minus-end of microtubules is mediated by a single motor, cytoplasmic Dynein (hereafter referred to as Dynein) (Reck-Peterson et al., 2018). The exact mechanism by which Dynein is able to transport a plethora of diverse cargo is unknown. Dynein is thought to engage numerous adaptors which serve to link cargo with the motor complex (Reck-Peterson et al., 2018). The identity of some of these adaptors are known, but given the diversity of cargo, it is likely that many more adaptors and regulators of Dynein exist.

An excellent model for examining the mechanism of Dynein-mediated transport is the egg chamber of *Drosophila melanogaster* (Goldman and Gonsalvez, 2017). The *Drosophila* egg chamber consists of fifteen nurse cells, a single oocyte, and layer of somatic follicle cells (Fig. 1). The nurse cells and oocyte are connected via structures known as ring-canals (Robinson and Cooley, 1996). Thus, these cells share a common cytoplasm. Within early-stage egg chambers (stages1 through 6), microtubules are organized such that minus-ends are enriched within the oocyte and plus-ends extend through the ring canals into the nurse cells (Fig. 1) (Riechmann and Ephrussi, 2001). As a consequence of this microtubule organization, cargoes transported by Dynein are typically enriched within the oocyte. Transport appears to be almost unidirectional, with most particles moving from the nurse cells into the oocyte (Lu et al., 2021). Consistent with a role for Dynein in mediating this transport, components of the motor, and Dynactin, an essential regulator of Dynein, are enriched within the oocyte of early stage egg chambers (Januschke et al., 2002; Li et al., 1994). The oocyte enrichment of Dynein requires Egalitarian (Egl). When Egl is depleted or mutated, Dynein transport into the oocyte is severely compromised (Goldman et al., 2021; Goldman et al., 2019; Navarro et al., 2004).

**Figure 1:**
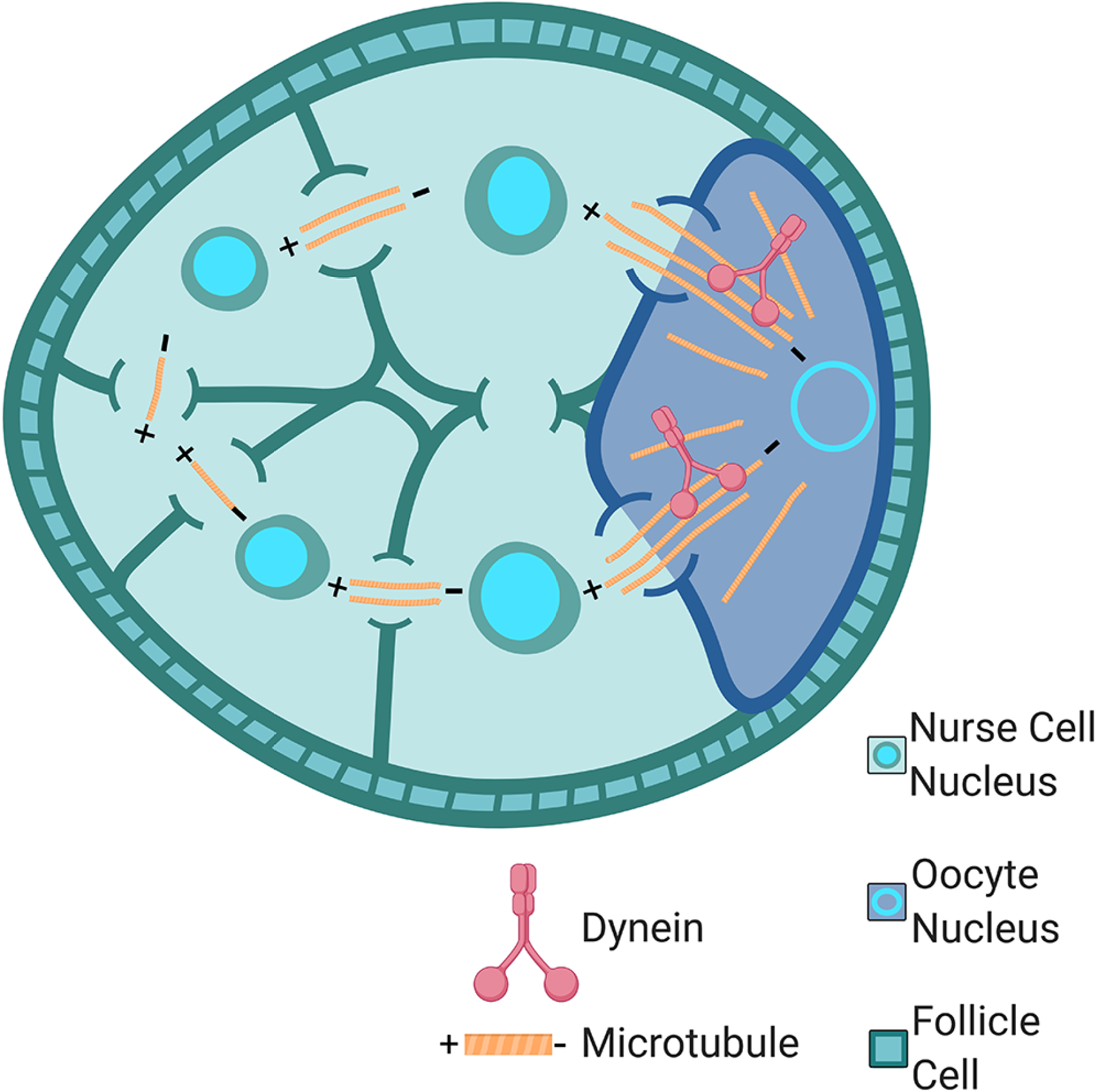
Illustration of a stage5 egg chamber. The nurse cells, oocyte, and somatic follicle cells are indicated. The organization of microtubules with their minus ends enriched within the oocyte is also indicated. Microtubules growing from the oocyte extend into the nurse cells via ring canals.

Egl is an RNA binding protein and is thought to link mRNA cargo with the Dynein motor. Once Egl binds to mRNA, it is able to interact with BicD, which in turn links this complex with Dynein (Dienstbier et al., 2009; Goldman et al., 2019; McClintock et al., 2018; Sladewski et al., 2018). As such, mRNAs that are known to be associated with Egl, are typically enriched within the oocyte (Goldman et al., 2021; Vazquez-Pianzola et al., 2017). However, in addition to mRNAs, numerous additional cargoes such as vesicles of the endoplasmic reticulum (ER), Golgi, the endocytic coat protein Clathrin, and lipid droplets are also oocyte enriched (Lee and Cooley, 2007; Lu et al., 2021; Roper, 2007; Vazquez-Pianzola et al., 2014). These cargoes are presumably transported into the oocyte in a Dynein-dependent manner. Given that the transport of Dynein into the oocyte depends upon Egl, we wondered whether Egl might also be involved in facilitating the transport of other types of cargo. In order to explore this topic, we chose to identify the Egl interactome.

Transport complexes containing Egl are likely to be highly dynamic and held together by weak and transient interactions. Thus, traditional biochemical strategies such as co-immunoprecipitation are not suitable for identifying the components of these complexes. For instance, although an Egl/BicD/Dynein complex can be assembled in vitro (McClintock et al., 2018; Sladewski et al., 2018), this complex cannot be isolated from fly lysates using a standard co-immunoprecipitation protocol. A weak interaction can be observed between Egl and Dynein if exogenous mRNA is added to the lysate. However, under native conditions, this complex appears to dissociate during purification (McClintock et al., 2018). We therefore used proximity biotin ligation to identify the Egl interactome. A newly described promiscuous biotin ligase termed TurboID (hereafter referred to as Trbo) was used in these experiments (Branon et al., 2018). We chose to use Trbo because unlike BioID, Trbo is active at 25°C (Branon et al., 2018), the optimal temperature for maintaining flies. Using this approach, we identified several previously-unknown interactors of Egl. These included components of P bodies and factors that were shown to interact with Dynein in mammalian cells. In addition, we developed and validated a nanobody based Trbo approach, which we refer to as GBP-Trbo. This approach enables the quick and easy interactome identification of GFP-tagged proteins. This tool is especially useful for the *Drosophila* community given the thousands of publicly available GFP tagged strains (Buszczak et al., 2007; Kelso et al., 2004; Nagarkar-Jaiswal et al., 2015; Sarov et al., 2016). Using GBP-Trbo, any lab wishing to perform an interactome analysis can do so with a single cross.

## RESULTS

### Egalitarian tagged with TurboID biotinylates interacting proteins

The goal of this study was to identify the interactome of Egalitarian (Egl), a cargo adaptor of the Dynein motor. The strategy we chose to pursue was in vivo proximity biotin ligation using TurboID (hereafter referred to as Trbo) (Branon et al., 2018). Egl was tagged at its C-terminal end with Trbo (Fig. 2A). This construct also contained a 3xFLAG tag and silent mutations within the coding region of *egl* that prevents its recognition by an *egl*-specific shRNA. This shRNA, when driven using a maternal Gal4 driver, is capable of depleting endogenous Egl within the female germline (Sanghavi et al., 2016). This strategy enables us to express Egl-Trbo in a background that is depleted of endogenous Egl. Egg chambers from flies expressing Egl-FLAG and Egl-Trbo were processed for immunofluorescence using a FLAG antibody. The egg chambers were also stained using Streptavidin conjugated with Alexa647 (Strep647). Streptavidin binds to biotin and therefore enables the visualization of in vivo biotinylated proteins. Both Egl-FLAG and Egl-Trbo were highly enriched in the oocyte, and also at the nurse cell nuage (Fig. 2B, C, arrow and arrow-head respectively). This localization pattern is identical to untagged Egl (Mach and Lehmann, 1997). Minimal Strep647 signal was observed in flies expressing Egl-FLAG (Fig. 2B’). By contrast, flies expressing Egl-Trbo produced robust Strep647 signal in a pattern that co-localized with Egl-Trbo (Fig. 2C’, C’’).

**Figure 2:**
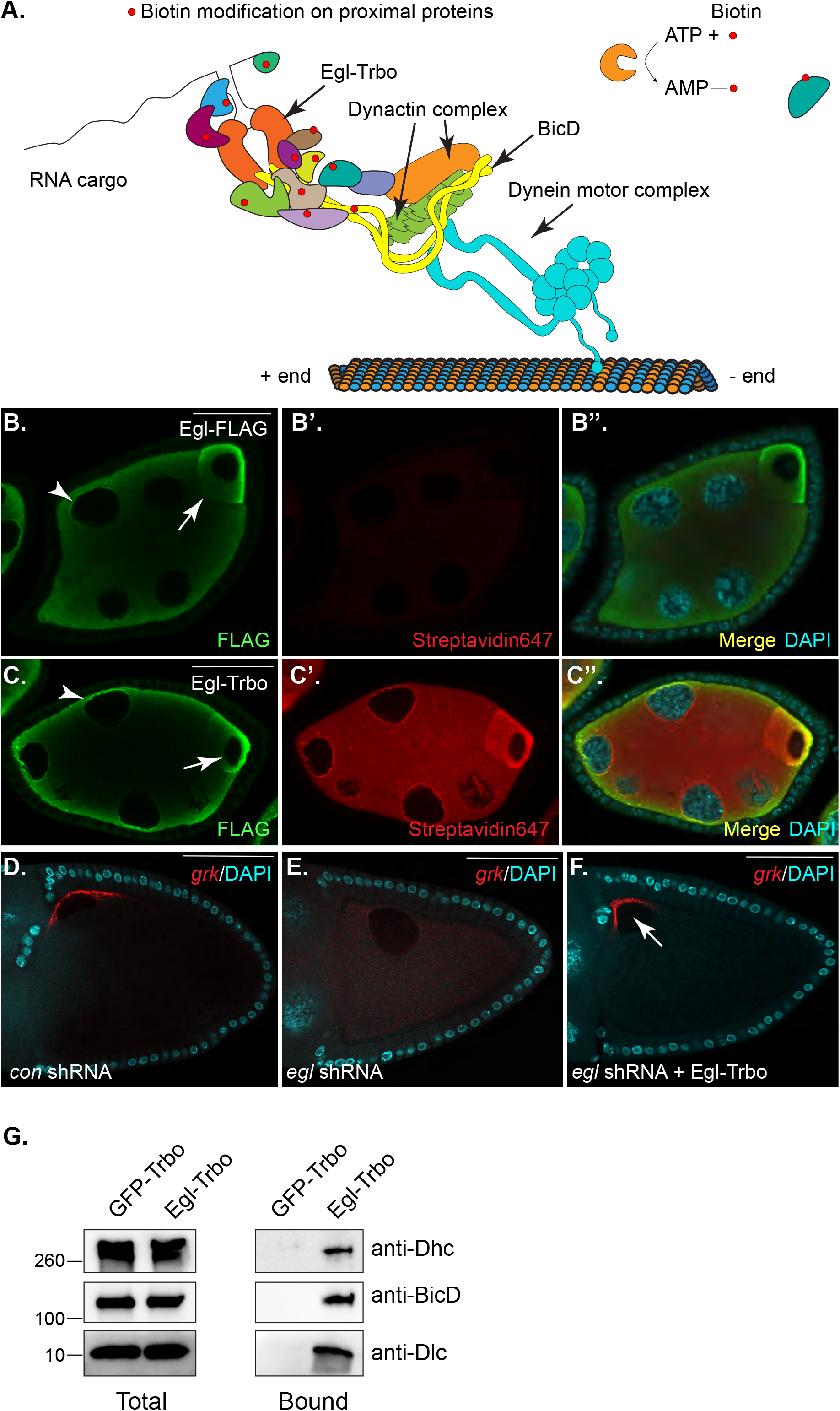
Egl-Tbro biotinylates interacting proteins. **(A)** Schematic of the Egl-BicD-Dynactin-Dynein complex. Egl-Trbo biotinylates proximal proteins (red circles). **(B-C)** Egg chambers expressing Egl-FLAG (B) or Egl-Trbo (C, which also contains a FLAG tag) were processed for immunofluorescence using a FLAG antibody (green). The samples were also processed using Streptavidin647 (red, B’, C’) and counterstained with DAPI (cyan). A merged image is shown (B’’, C’’). **(D-F)** Ovaries were dissected from the indicated genotypes and processed for in situ hybridization using probes against *grk* (red). The egg chambers were counterstained with DAPI to reveal nuclei (cyan). The arrow indicates the dorsal-anterior localization of *grk* mRNA. **(G)** Ovarian lysates were prepared from strains expressing GFP-Trbo or Egl-Trbo. Biotinylated proteins were purified and analyzed by western blotting using the indicated antibodies. The scale bar in B and C is 20 microns and the scale bar in D-F is 50 microns.

Next, we determined whether Egl-Trbo is functional. Depletion of Egl results in the delocalization of *grk* mRNA (Fig. 2D, E) (Sanghavi et al., 2016). In addition, embryos from Egl-depleted mothers do not survive (Sanghavi et al., 2016). Expression of Egl-Trbo in this background restored *grk* mRNA localization (Fig. 2F, arrow). Furthermore, embryos from these females were completely viable (data not shown). Thus, Egl-Trbo is functional and able to compensate for the loss of endogenous Egl. We next purified biotinylated proteins from ovarian lysate expressing either GFP-Trbo or Egl-Trbo using Streptavidin conjugated beads. As expected, BicD and Dynein light chain (Dlc), known interacting partners of Egl (Mach and Lehmann, 1997; Navarro et al., 2004), were highly enriched in the Egl-Trbo pellet in comparison to the control (Fig. 2G). Dynein heavy chain (Dhc), the motor subunit of Dynein, was also enriched in the Egl pellet (Fig. 2G). Based on these results, we conclude that Egl-Trbo is localizing as expected, is functional, and is biotinylating proximal proteins.

### Defining the Egl interactome

In order to identify the Egl interactome, ovaries were dissected from flies expressing either GFP-Trbo or Egl-Trbo. Lysates were prepared and the biotinylated proteins were purified using Streptavidin conjugated beads. The beads were extensively washed, the bound proteins were digested with trypsin, and the samples were analyzed using mass-spectrometry. The entire experiment was done in triplicate. Proteins enriched at least two-fold in the Egl pellet with a p value of at least 0.05 were considered specific interacting partners (Fig. 3A, Supplemental table 1).

**Figure 3:**
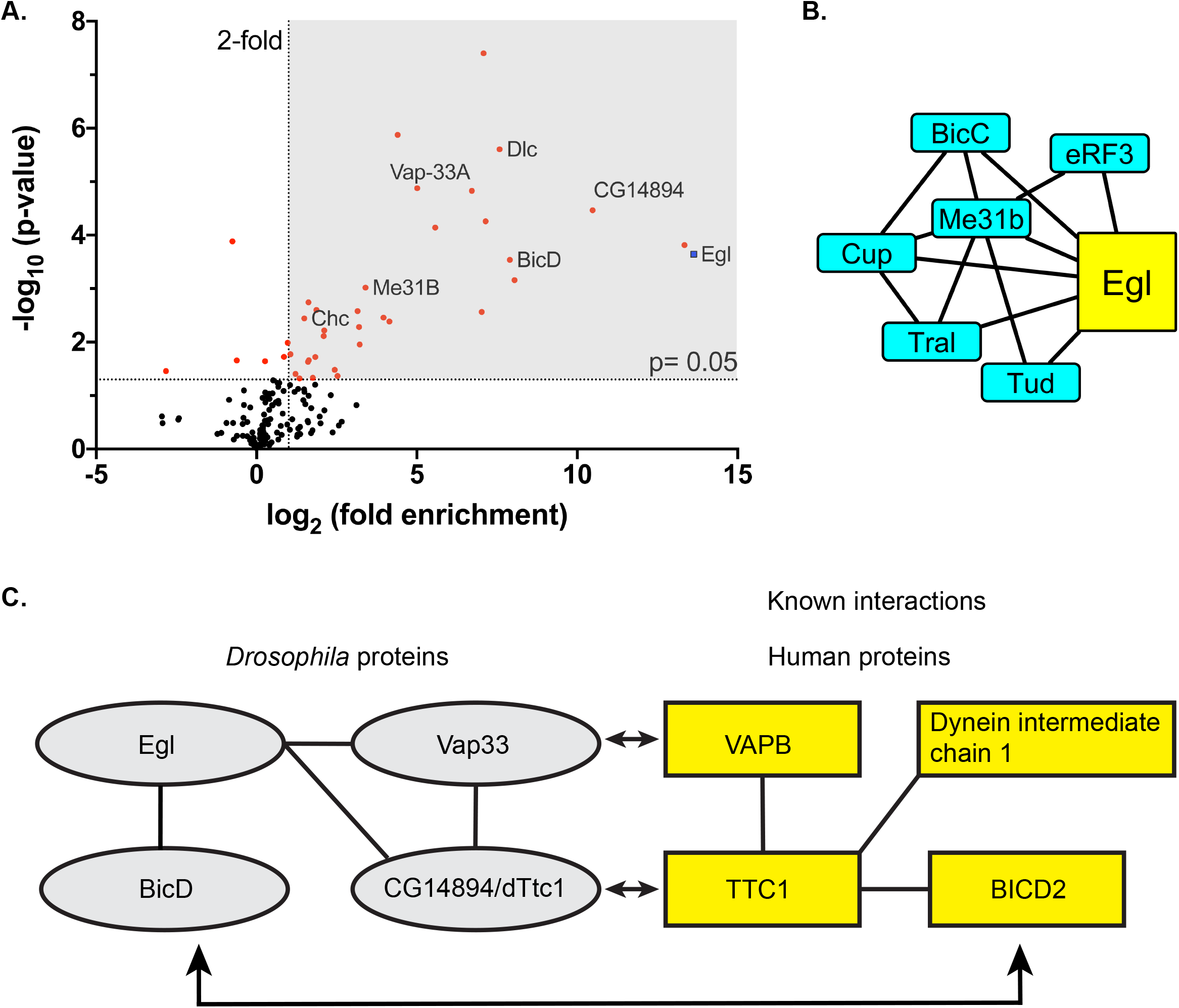
The Egl interactome. **(A)** A volcano plot depicting the Egl interactome. A line demarcating 2 fold enrichment and a p value of 0.05 is shown. **(B)** A protein-protein interaction network of P body components that were identified in the Egl interactome. **(C)** A schematic illustrating the interactions between *Drosophila* and human proteins. The interaction between Egl, Vap33 and dTtc1 were identified in this study. The interactions between TTC1 and BICD2 and Dynein intermediate chain, as well as between TTC1 and VAPB were previously described (citations within the text).

Egl has been shown to exist as a dimer (Goldman et al., 2019; Mach and Lehmann, 1997). Thus, Egl-Trbo should self-associate and result in cross-biotinylation. Consistent with this notion, peptides corresponding to Egl were highly enriched in the Egl-Trbo interactome (Fig. 3A). In addition, BicD and Dlc, known interacting partners of Egl (Mach and Lehmann, 1997; Navarro et al., 2004), were highly enriched in the Egl interactome (Fig. 3A). Clathin heavy chain (Chc) was also enriched in the Egl interactome (Fig. 3A). Although, Chc has not yet been shown to interact with Egl, it is present in a complex with BicD in the female germline and in neurons (Li et al., 2010; Vazquez-Pianzola et al., 2014). Based on our findings, it is likely this Chc/BicD complex in oocytes also contains Egl.

Surprisingly, although Dhc could be detected in the Egl-Trbo pellet by western blot (Fig. 2G), it was not detected in the interactome using mass-spectrometry. The reason for this might have to do with the higher sensitivity of western blotting, combined with the limited biotinylation radius of Trbo, estimated to be between around 35nm (Kim et al., 2014; May et al., 2020). It has been previously demonstrated that components of the nuclear pore complex or Dynein that are further away from the tagged protein are not as efficiently biotinylated in comparison to components that are closer. This results in these more distal components being inefficiently purified and thus under-represented by mass-spectrometry analysis (Kim et al., 2014; Kim et al., 2016; Redwine et al., 2017).

### P body components are enriched in the Egl-Trbo pellet

The Egl interactome revealed several novel interactions with known P body components (Fig. 3B). P bodies are cytoplasmic aggregates of protein/RNA complexes involved in regulating the translation and stability of mRNAs (Youn et al., 2019). In addition to their potential interaction with Egl, several of these proteins have been previously shown to interact with each other (Fig. 3B). We focused our initial studies on the P body component, Me31b. We repeated the purification on a smaller scale and analyzed the pellets by western blotting. Consistent with the proteomics result, BicD and Me31b were highly enriched in the Egl-Trbo pellet in comparison to the control (Fig. 4A).

**Figure 4:**
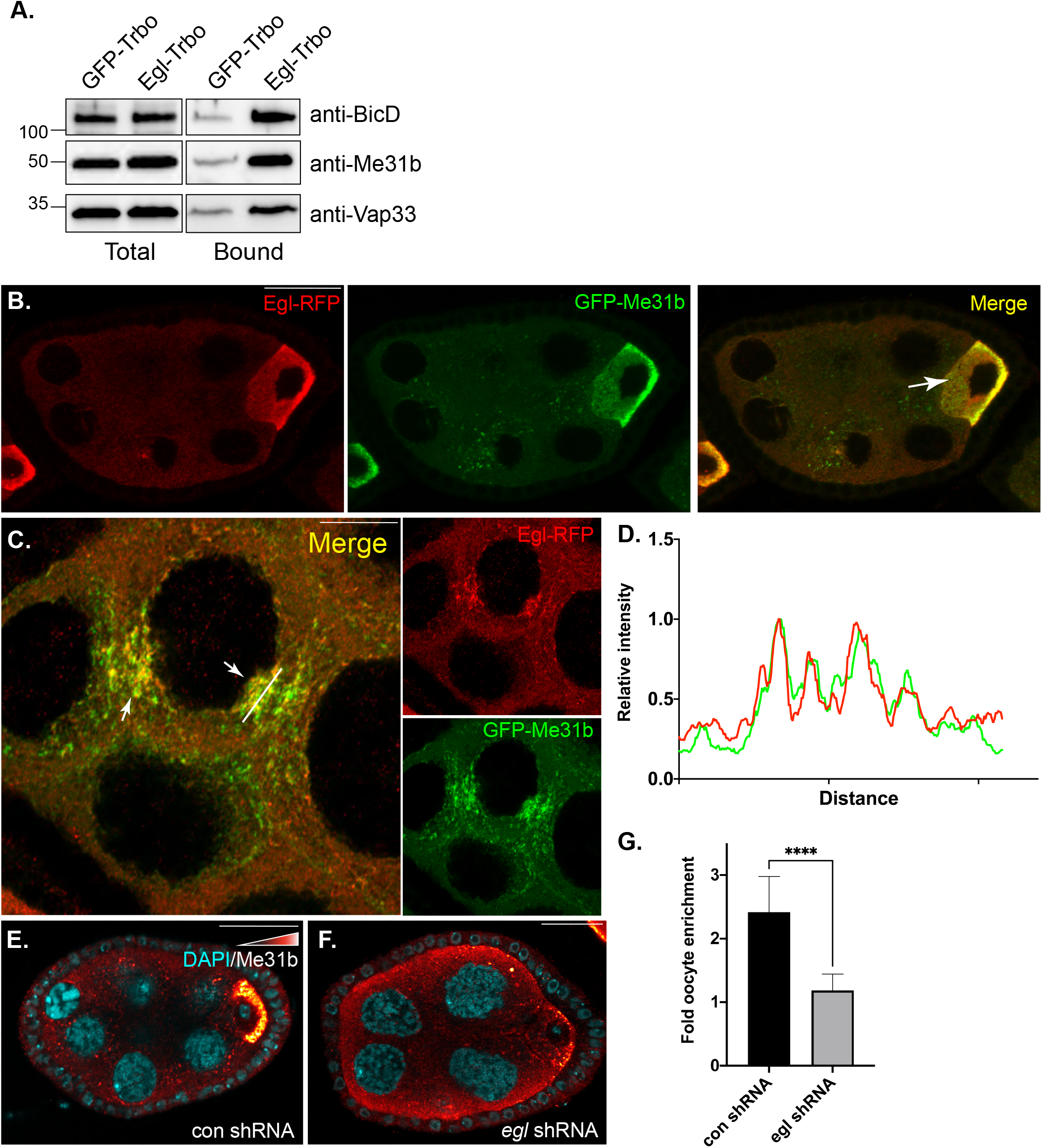
The Egl-P body interaction. **(A)** Lysates from the indicated genotypes were prepared and the biotinylated proteins were purified. Bound proteins were analyzed by western blotting using the indicated antibody. BicD, Me31b and Vap33 specifically precipitate in lysates expressing Egl-Trbo. **(B-D)** A strain expressing Egl-RFP and GFP-Me31b was processed for immunofluorescence using an antibody against RFP. Both proteins are highly enriched within the oocyte (B, arrow). Within the nurse cell cytoplasm, Egl-RFP was often detected co-localizing with or adjacent to Me31b foci (C, arrows). Panel D shows intensity values of Egl-RFP and GFP-Me31b in the region of the line depicted in panel C. **(E-G)** Egg chamber expressing a control shRNA (*eb1* shRNA) (E) or an shRNA against *egl* (F) were fixed and processed using an antibody against Me31b. The signal for Me31b is depicted using a red to white look up table (LUT), where red indicates pixels of low intensity and white indicates pixels of high intensity. The oocyte enrichment of Me31b was calculated and plotted in panel G. Efficient enrichment of Me31b in the oocyte requires Egl. An unpaired *t* test was used for this analysis; ****p<0.0001.

In addition to their biochemical interaction, Egl and Me31b extensively co-localized. Both proteins were highly enriched in the oocyte (Fig. 4B, arrow), and in the nurse cell cytoplasm, Egl foci were often adjacent to or co-localized with P bodies (Fig. 4C, D). We next determined whether the localization of Me31b was dependent upon Egl. The oocyte enrichment of Me31b was significantly reduced in strains that were depleted of Egl (Fig.4 E, F, G). P bodies were still detected in these egg chambers (Supplemental Fig.1A). However, a qualitative difference was noted. In controls, the signal for Me31b was highly enriched in P bodies with minimal diffuse cytoplasmic staining. By contrast, in Egl depleted flies, although P bodies were still detected, there was considerably more diffuse signal for Me31b (Supplemental Fig.1A’). Thus, Egl appears to be primarily required for efficient transport of Me31b into the oocyte, with perhaps a secondary function in facilitating the enrichment of Me31b into P bodies.

Results from our lab and others indicate that loss of Me31b and Egl produce similar phenotypes; in both cases *osk* mRNA is delocalized and inappropriately translated (Nakamura et al., 2001; Sanghavi et al., 2016). These findings, combined with our identification of several P body components in the Egl interactome, suggest a potential functional relationship between Egl and P bodies. Given that most localized mRNAs are also regulated at the level of translation (Lasko, 2012), the Egl-P body interaction might suggest that localized mRNAs are first trafficked through P bodies. This would enable the localization of these mRNAs to be tightly coupled to their translational regulation.

### Egl interacts with Vap33 and dTtc1

Two additional proteins in the Egl interactome were of particular interest to us: CG14894 and Vap33 (Fig. 3A). CG14894, an uncharacterized gene, is the *Drosophila* ortholog of human TTC1 (Tetratricopeptide Repeat Domain 1). We will therefore refer to the *Drosophila* protein as dTtc1 (Fig. 3C). We were able to validate the interaction between Egl and Vap33 (Fig. 4A). However, at present, an antibody against dTtc1 is not available. Thus, we are not able to validate the Egl-dTtc1 interaction by western blotting.

The Reck-Peterson lab recently performed an interactome study of the Dynein motor. In this study, several components of the motor as well as cargo adaptors were tagged with BioID and expressed in HEK293T cells (human embryonic kidney epithelial cells) (Redwine et al., 2017). Interestingly, TTC1 was enriched in the pellets of BioID tagged BicD2 and Dynein intermediate chain (Dic) (Redwine et al., 2017). BicD2 is the human ortholog of *Drosophila* BicD, and Dic is a core component of the Dynein motor (Hoogenraad and Akhmanova, 2016; Reck-Peterson et al., 2018). Consistent with the proteomics results of Redwine et al., we observed a specific interaction between Trbo-TTC1 and BicD2 in HeLa cells (Supplemental Fig.1B). In addition to its association with the Dynein motor, TTC1 also interacts with VAPB, the human ortholog of fly Vap33 (Lotz et al., 2008; Taipale et al., 2014) (Fig. 3C). Thus, these interactions between components of the Dynein motor, TTC1, and VAPB/Vap33 appear to be conserved between fly oocytes and mammalian cells.

Studies in mammalian cells indicate that VAPB is an endoplasmic reticulum (ER)-resident protein and is often detected at sites at which ER membranes contact organelles such as Golgi, mitochondria, endosomes, lysosomes and peroxisomes (Dudas et al., 2021; Kamemura and Chihara, 2019). Mutations in VAPB are associated with Amyotrophic lateral sclerosis (ALS) and a form of Spinal muscular atrophy (Nishimura et al., 2004). In flies, Vap33 is an essential gene, and most studies to date have focused on the role of this protein in neuronal cells. Whether or not Vap33 also has a role in the germline is unknown.

In the germarium, Vap33 and Calnexin (Cnx99A), an ER integral membrane protein extensively colocalized in a perinuclear region (Supplemental Fig.1C). In stage2 egg chambers expressing Egl-GFP, all three proteins co-localized around the perinuclear region surrounding nurse cell nuclei (Fig. 5A, B). The localization of Vap33 was dynamic and in stage5 egg chambers, Egl-GFP and ER vesicles could still be detected around nurse cell nuclei, likely at the nuage (Fig. 5C). In these egg chambers, Vap33 was not nuage-enriched. However, all three proteins were highly enriched within the oocyte of these stage5 egg chambers (Fig. 5C, arrow).

**Figure 5:**
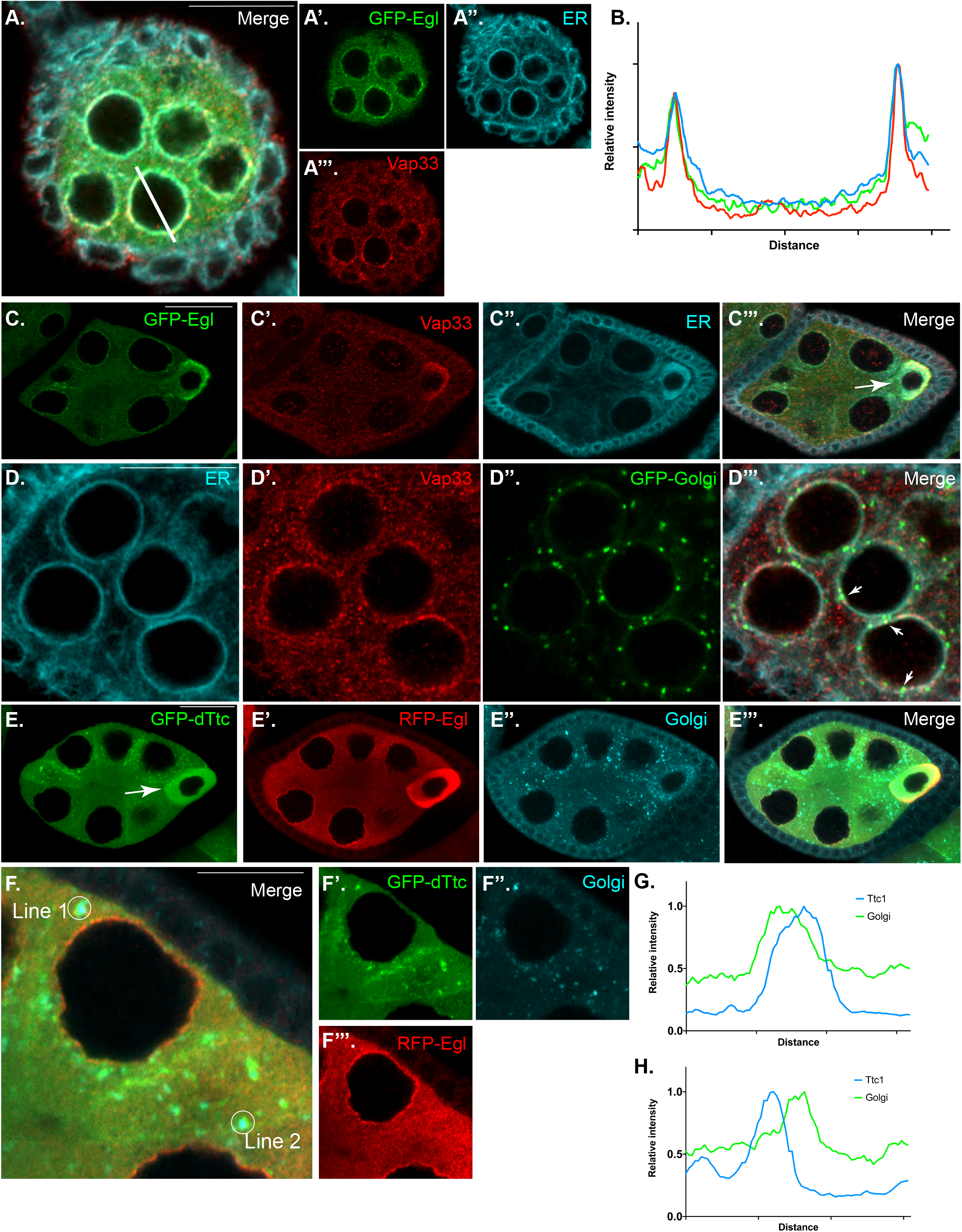
Localization of Vap33 and dTtc1 in the female germline. **(A-C)** Egg chambers from flies expressing Egl-GFP (green) were fixed and processed for immunofluorescence using antibodies against Vap33 (Red) and Cnx99A (cyan), an ER vesicle maker. Panel A depicts the localization of these proteins in a stage2 egg chamber and panel B shows the intensity profile of the line depicted in panel A. Panel C shows the localization of these proteins in a stage5 egg chamber. Egl, Vap33 and ER vesicle are highly enriched in the oocyte (arrow). (**D)** A strain expressing a GFP-tagged Golgi maker (Galactosyltransferase-GFP, creen) was processed using antibodies against Vap33 (red) and Cnx99A (cyan). Vap33 could often be detected at sites of co-localization between Golgi and ER (arrows). **(E-H)** A strain co-expressing GFP-dTtc1 (green) and Egl-RFP (red) was fixed and processed using an antibody against a Golgi marker (cyan). Egl-RFP and GFP-dTtc1 were highly enriched in the oocyte, whereas Golgi vesicles were moderately oocyte-enriched (E, arrow). Within the nurse cells, GFP-dTtc1 foci were often detected co-localizing with or adjacent to Golgi foci (F). Representative line scans are shown in panels G (line1) and H (line 2). However, in order to avoid obscuring the localization of these proteins, the lines used for quantification are represented as open circles. The scale bar in F is 10 microns. In the remaining images, the scale bar is 20 microns.

Functional studies indicate that Vap33 is required for an ER to Golgi trafficking step (Mao et al., 2019). We therefore examined the localization of Vap33 and ER vesicles in egg chambers expressing a GFP tagged Golgi marker. Consistent with a function for Vap33 in mediating ER to Golgi transport, Vap33 could often be detected at potential sites of ER-Golgi contact (Fig. 5D, arrows).

Very little is known regarding the function of human TTC1 or its *Drosophila* ortholog. At present, an antibody against dTtc1 is not available. We therefore generated a strain that expresses GFP-dTtc1 in the female germline. Like Vap33 and Egl, GFP-Ttc1 was also oocyte enriched (Fig. 5E, arrow). However, unlike Vap33, GFP-dTtc1 did not display a nuage localization pattern. Numerous foci of GFP-dTtc1 were observed close to the perinuclear region surrounding nurse cell nuclei. Often, these foci of GFP-dTtc1 co-localized with or were adjacent to a marker for Golgi vesicles (Fig. 5F, G, H).

The oocyte enrichment of Vap33 and dTtc1 suggests that they might be transported into the oocyte in a Egl/Dynein dependent manner. We therefore examined the localization of these proteins in Egl depleted egg chambers. In contrast to egg chambers expressing a control shRNA, the oocyte enrichment of both Vap33 and GFP-dTtc1 were reduced in egg chambers expressing the *egl* shRNA (Fig. 6A-D, I). A similar phenotype was observed for ER vesicles (Fig. 6E, F). The distribution of Golgi, however, was relatively unaffected by the depletion of Egl (Fig. 6G, H). Based on these observations, we conclude that efficient transport of Vap33, dTtc1 and vesicles of ER into the oocyte requires Egl.

**Figure 6:**
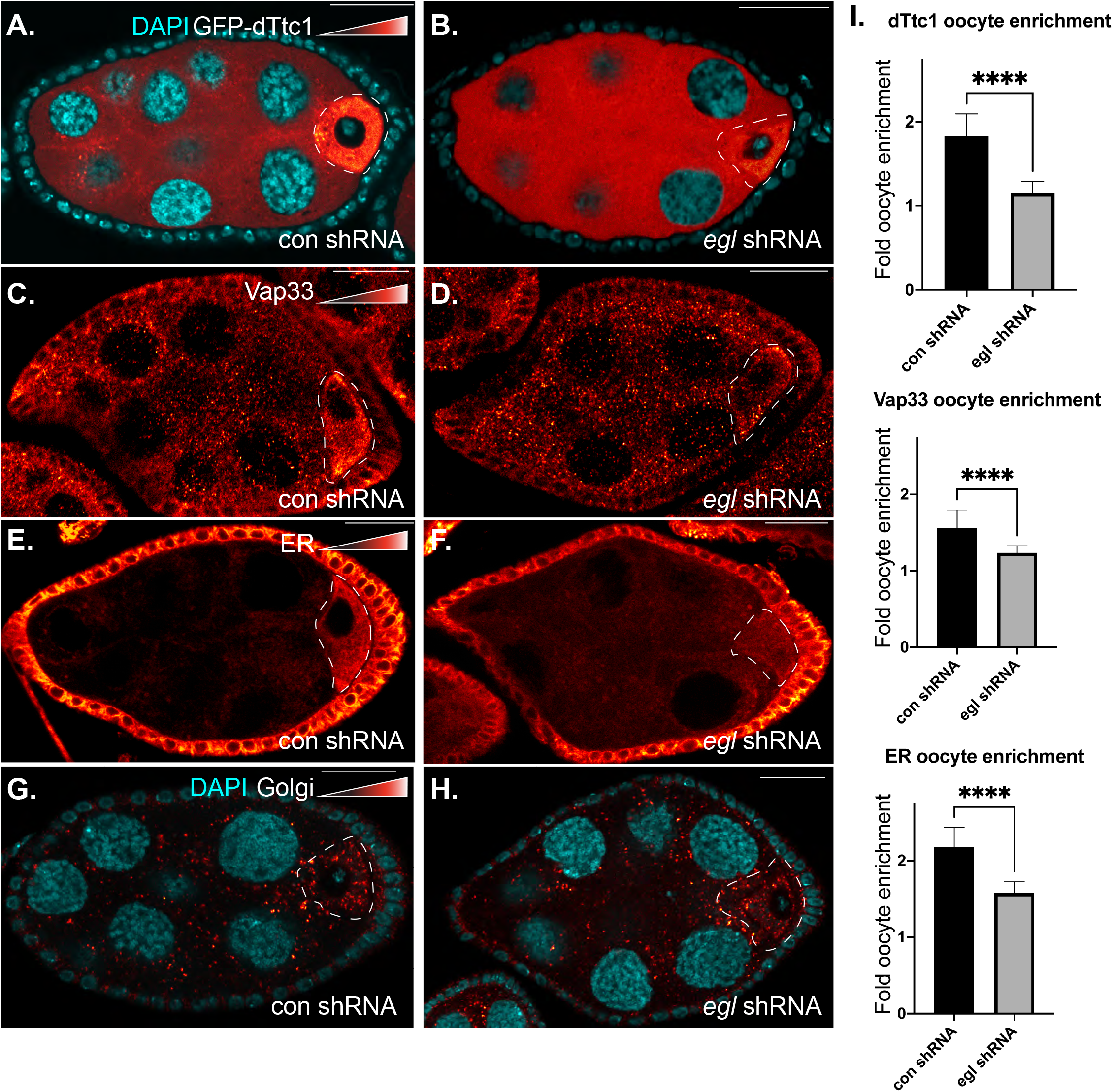
Oocyte enrichment of dTtc1, Vap33 and ER vesicles requires Egl. **(A-H)** Egg chambers from flies expressing GFP-dTtc1 and a control shRNA (A) or an shRNA against *egl* (B) were fixed, DAPI stained, and imaged. Egg chambers expressing the same control shRNA (C, E, G) or an shRNA against *egl* (D, F, H) were fixed and processed using antibodies against Vap33 (C, D), ER vesicles (E, F) and Golgi vesicles (G, H). Signal for these proteins is represented using the red to white LUT. Oocyte enrichment of dTtc1, Vap33 and ER were dependent upon Egl, whereas the modest oocyte-enrichment of Golgi vesicles was not Egl-dependent. The oocyte is indicated by dashed lines. Quantification of this data is shown in panel I. Unpaired *t* tests were used for these analyses; ****p<0.0001. The scale bar is 20 microns.

### A GFP-nanobody approach to define interactomes

Why might dTtc1 associate with Egl? As noted previously, TTC1 is present in the Dynein interactome in mammalian cells (Redwine et al., 2017). Both fly and human Ttc1 contain three tetratricopeptide repeats (TPR), motifs that often mediate protein-protein interaction (Zeytuni and Zarivach, 2012). Kinesin light chain, a cargo adaptor of the plus-end directed Kinesin motor, contains TRP repeats with which it interacts with cargo (Cockburn et al., 2018; Hammond et al., 2008; Pernigo et al., 2013; Zhu et al., 2012). We thus hypothesized that dTtc1, and its human ortholog, function as cargo adaptors for Dynein. We therefore sought to determine the interactome of dTtc1.

One approach for determining the dTtc1 interactome would be to fuse it directly to Trbo. However, given that we were in possession of a strain in which dTtc1 was tagged with GFP, we decided to develop a nanoboby based approach. A nanobody is a small protein, usually 15kD or less, that consists of the antigen-recognition domain of a single chain antibody (de Beer and Giepmans, 2020). We fused the GFP nanobody, referred to as the GFP binding protein (GBP), to Trbo. We refer to this as GBP-Trbo. In theory, GBP-Trbo should bind to any GFP tagged protein in vivo and result in the biotinylation of proximal proteins. We decided to develop this tool due to the abundant public availability of thousands of GFP tagged fly strains (Buszczak et al., 2007; Kelso et al., 2004; Nagarkar-Jaiswal et al., 2015; Sarov et al., 2016). In addition to these easily available strains, most labs routinely tag their protein of interest (POI) with GFP in order to examine its localization in vivo. Thus, GBP-Trbo could be a very useful tool for the *Drosophila* community and would enable any lab to easily define the interactome of a GFP tagged protein.

In order to validate this approach, we co-expressed GBP-Trbo along with GFP. GFP was diffusely localized in the egg chamber. The same localization pattern was observed using Strep647, a reporter for biotinylated proteins (Fig. 7A). We next co-expressed GBP-Trbo along with GFP-dTtc1. As shown in a previous figure, GFP-dTtc1 is enriched within the oocyte and also present in numerous foci in the nurse cell cytoplasm (Fig. 7B). The Strep647 pattern co-localized with GFP-dTtc1 (Fig. 7B, C). A similar co-localization was observed in strains co-expressing GBP-Trbo and GFP-Me31b (Fig. 7D, E). Of note, the GFP-Me31b strain used in this experiment was generated in a Flytrap screen (Buszczak et al., 2007) and was obtained from the Bloomington stock center. GFP is inserted at the endogenous *me31b* locus in this strain. Thus, the localization of GFP-Me31b represents its endogenous expression pattern.

**Figure 7:**
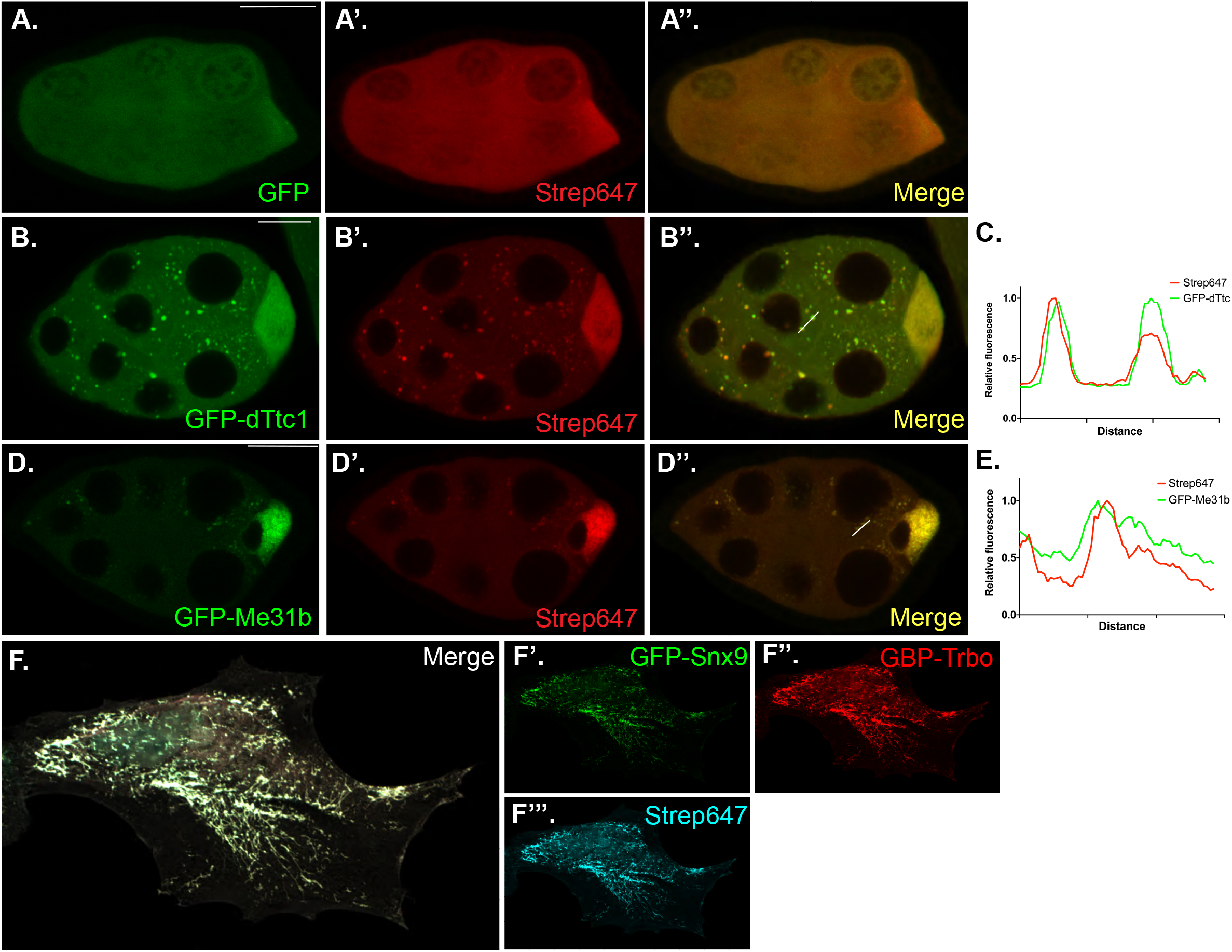
GBP-Trbo binds to GFP tagged proteins and biotinylates proximal proteins. **(A)** Egg chambers from flies co-expressing GFP (green) and GBP-Trbo were fixed and processed using Streptavidin674 (red). A merged image is also shown. **(B-C)** Egg chambers from flies expressing GFP-dTtc1 (green) were fixed and processed using Streptavidin674 (red). Streptavidin and GFP signals co-localize (B’’). A line scan of intensity values is shown in panel C. **(D-E)** Egg chambers from flies expressing GFP-Me31b (green) were fixed and processed for Streptavidin674 (red). As with GFP-dTtc1, the Streptavidin and GFP signals co-localize (D’’). A line scan of intensity values is shown in panel E. **(F)** HeLa cells were transfected with plasmids expressing GFP-Snx9 (green) and GBP-Trbo (red). The cells were fixed and processed using a V5 antibody to detect GBP-Trbo. The cells were also processed using Streptavidin674 (cyan). Individual channels and a merged image is shown.

In addition to *Drosophila* egg chambers, we found that this approach also works quite well in mammalian cells. HeLa cells were co-transfected with GBP-Trbo and GFP-Snx9. Snx9 over-expression in mammalian cells results in the formation of long membrane tubules (Shin et al., 2008). GBP-Trbo and the resulting Strep647 signal co-localized on these membrane tubules (Fig. 7F). Collectively, these results suggest that GBP-Trbo is a valid approach for tethering a biotin ligase to a GFP-tagged POI.

### Defining the Me31b interactome using GBP-Trbo

In order to determine whether the GBP-Trbo approach would work for interactome analysis, we used the afore mentioned GFP-Me31b strain. Me31b is a good candidate for validating this approach because it has a well-documented function in the female germline (Davidson et al., 2016; Gotze et al., 2017; Nakamura et al., 2001). In addition, several of its interacting partners have been previously identified using either low-throughput co-immunoprecipitation experiments or using chemical cross linkers followed by immunoprecipitation and mass spectrometry (DeHaan et al., 2017; Liu et al., 2011; McCambridge et al., 2020; Nakamura et al., 2004; Wilhelm et al., 2005). Ovaries were dissected from flies expressing either GFP alone or GFP-Me31b. Biotinylated proteins were purified and identified using mass spectrometry. The entire experiment was done in triplicate. Proteins enriched at least two-fold in the GFP-Me31b pellet with a p value of at least 0.05 were considered specific interacting partners (Fig. 8A, Supplemental table2).

**Figure 8:**
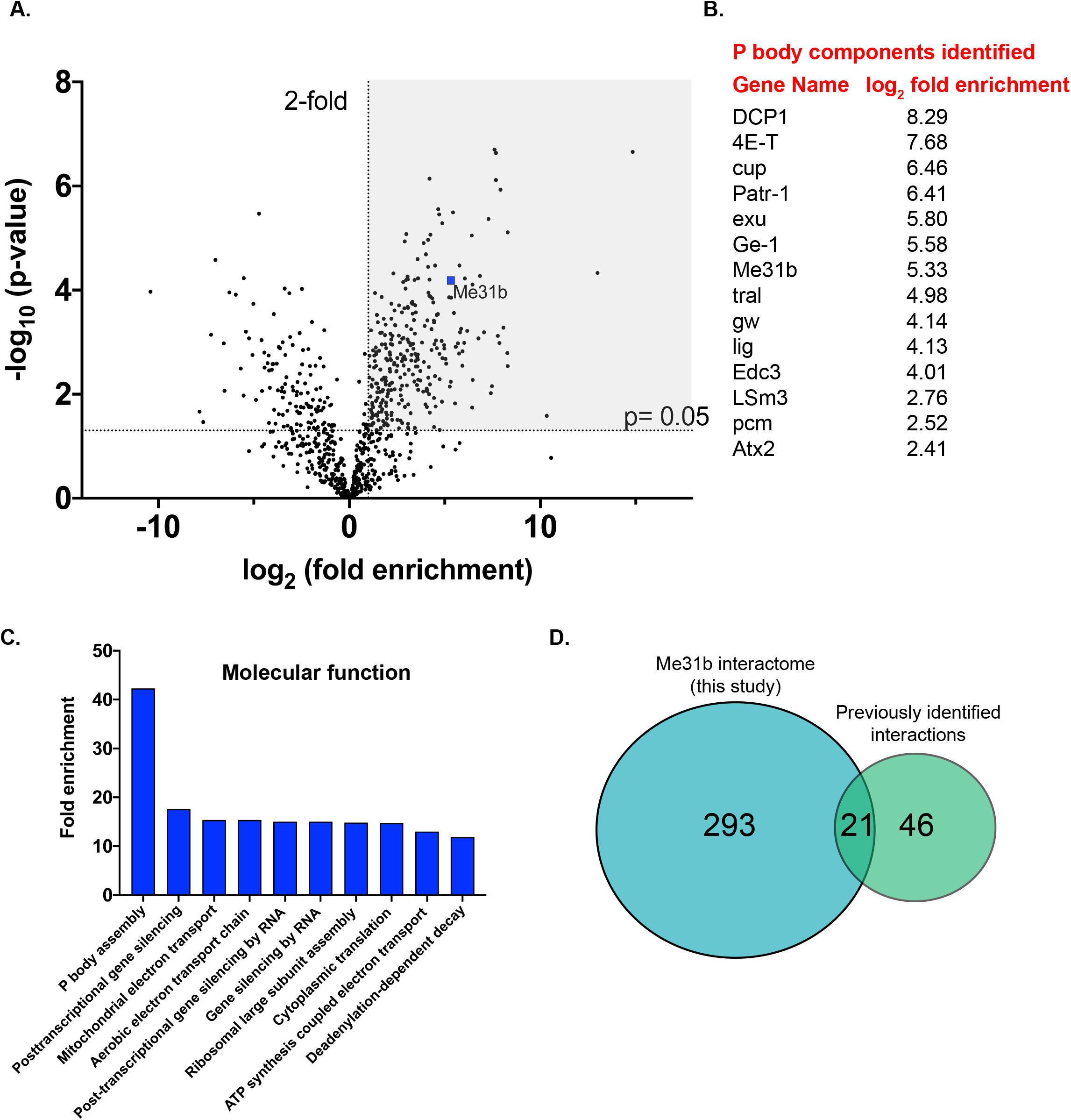
The Me31b interactome. **(A)** A volcano plot depicting the Me31b interactome. A line indicating 2 fold enrichment and a p value of 0.05 is shown. **(B)** This list shows the P body components enriched in the Me31b interactome. **(C)** A GO analysis of molecular function revealed an enrichment of proteins with known roles in post-transcriptional gene expression. **(D)** A comparison between the number of protein interactions that were previously described for Me31b and the number of interactions obtained in our study.

As expected, Me31b was highly enriched in the interactome. In addition, out of a total of 39 predicted P body protein, 14 (including Me31b) were identified in the Me31b interactome (Fig. 8B). Consistent with the known function of Me31b, a GO analysis of molecular function revealed that the interactome contained an enrichment of factors that have a role in “P body assembly” and in various aspects of post-transcriptional gene expression (Fig. 8C). From previous studies, a total of 46 unique protein interactions have been described for Me31b. Twenty one of these interactions were also present in our data set (Supplemental table 3). Importantly, our analysis revealed a total of 293 unique protein interaction candidates for Me31b (including Egl), the vast majority of which were previously unknown (Fig. 8D). These results indicate that GBP-Trbo is a valid and powerful tool for defining the interactome of GFP-tagged proteins.

### The dTtc1 interactome

We next used a similar approach to determine the interactome of GFP-dTtc1. Flies expressing GFP alone were used as a control. As with our previous analysis, we considered proteins enriched at least two-fold in the GFP-dTtc1 pellet with a p value of at least 0.05 as specific interacting partners. This represents 175 unique interactions (Supplemental table 4). Not surprisingly, dTtc1 was highly enriched in the interactome (Fig. 9A). In addition, Vap33 was also highly enriched (Fig. 9A). Thus, like their mammalian counterparts, fly Vap33 and dTtc1 are also present in a complex.

**Figure 9:**
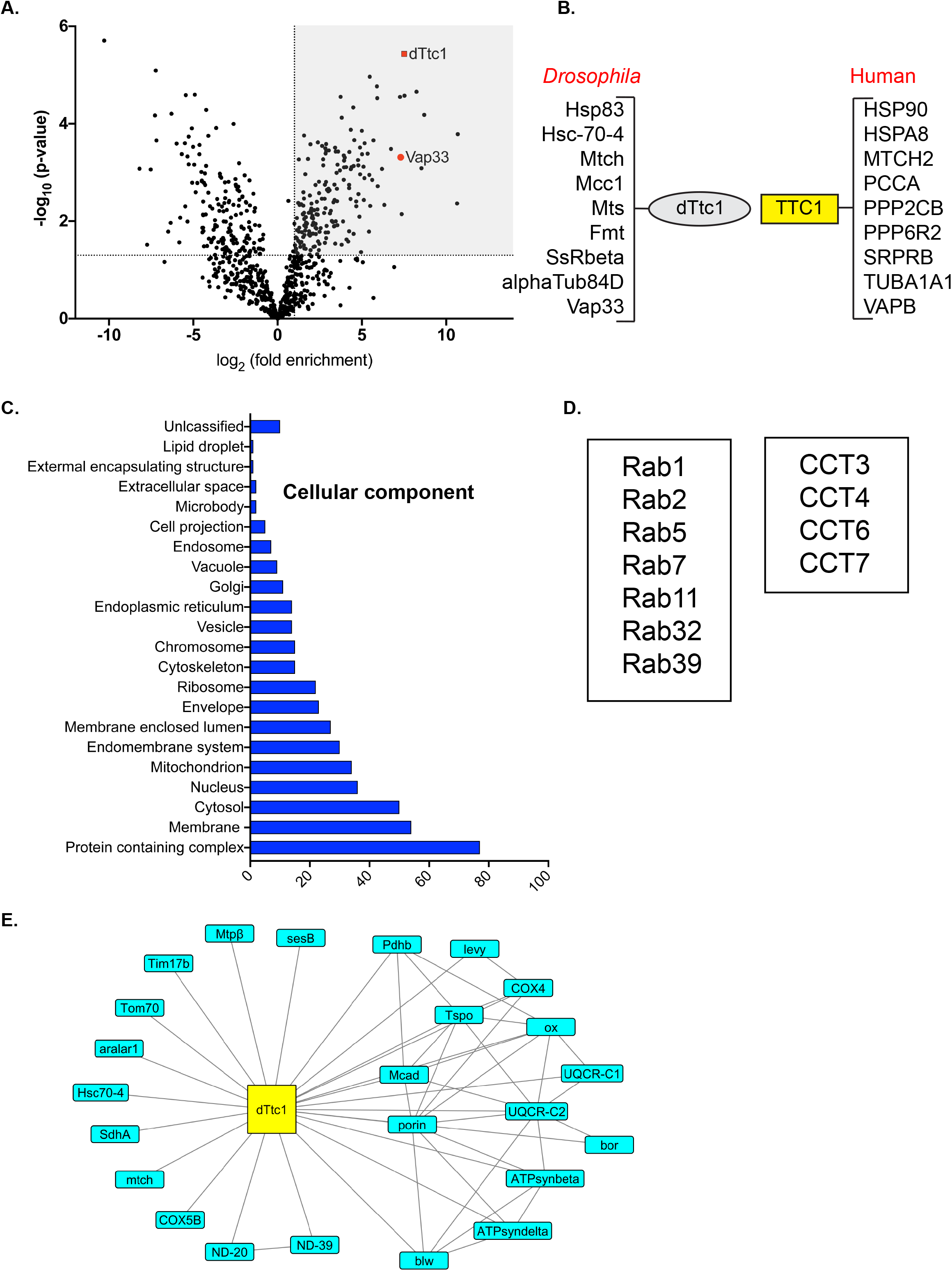
The dTtc1 interactome. **(A)** A volcano plot depicting the dTtc1 interactome. The layout of this plot is similar to the previous interactomes. **(B)** This list shows interactions between *Drosophila* dTtc1 and fly proteins that were also detected between human TTC1 and corresponding human orthologs. **(C)** A cellular compartment GO analysis revealed an enrichment of proteins that localize to the endomembrane system and other vesicular structures. **(D)** Several proteins of the Rab family and 4 out of 8 components of the CCT Chaperonin complex were found to interact with dTtc1. **(E)** dTtc1 interacts with numerous proteins that localize to mitochondria. Several of these proteins were previously shown to interact with each other.

Further confidence in the dTtc1 interactome comes from the observation of shared interactions between oocytes and mammalian cells. Although a proteomics screen has not been conducted in mammalian cells using TTC1 as the bait, TTC1 has been observed in the interactome of other proteins. In particular, 9 interactions that have been described for human TTC1 were also found as specific interacting partners of *Drosophila* dTtc1 (Fig. 9B). Within this list, the HSP90/Hsp83 interaction is noteworthy because VAPB, TTC1 and HSP90 are present in a complex in mammalian cells (Lotz et al., 2008). Our results suggests that this trimeric complex also forms in fly egg chambers.

A cellular component GO analysis of the dTtc1 interactome revealed an enrichment of terms such as “endomembrane system”, “cytoskeleton”, “endoplasmic reticulum”, “Golgi”, and “endosomes” (Fig. 9C). Consistent with this finding and a potential role for dTtc1 in vesicle trafficking, several Rab proteins were identified in the dTtc1 interactome (Fig. 9D). Rab proteins are small GTPases involved in regulating various aspects of vesicle trafficking (Wandinger-Ness and Zerial, 2014). Another interesting group of proteins in the interactome corresponds to factors that have a role in protein folding. The highly conserved CCT Chaperonin complex consists of 8 subunits, 4 of which were identified as specific interacting partners of dTtc1 (Fig. 9D). The main folding substrates of the CCT complex are actin and tubulin (Grantham, 2020). In addition, monomeric CCT4 has been shown to interact with the p150/Glued component of the Dynactin complex (Echbarthi et al., 2018). Lastly, several proteins within the dTtc1 interactome are predicted to localize to the mitochondrion (Fig. 9C, E). Interestingly, several of these proteins have been previously found to interact with each other (Fig. 9E). Thus, it is likely that dTtc1 is interacting with specific complexes of mitochondrial proteins.

In conclusion, dTtc1 interacts with a wide range of proteins across different cellular compartments. Based on the observation that dTtc1 is present is a complex with Egl (Fig. 3A) and with the Dynein motor in mammalian cells (Redwine et al., 2017), it is possible that at least some of the proteins will be linked to Dynein via their interaction with dTtc1. Testing this prediction will be the focus of a future study.

## DISCUSSION

### The Egl interactome

In this study, we used proximity biotin ligation to identify the interactome of Egl, a protein that has been shown to link mRNA with Dynein (Dienstbier et al., 2009). One group of proteins enriched in the Egl interactome were P body components. mRNAs that are localized are often translationally repressed until they are delivered to their cellular destination. This enables tight spatial restriction of the corresponding protein (Lasko, 2012). Given that P bodies have a well-documented role in regulating translation (Youn et al., 2019), it is possible that mRNAs localized in an Egl/Dynein-dependent manner are first trafficked through P bodies. This would enable localization and translational to be tightly linked. Consistent with this notion, several proteins that localize to P bodies have roles in mRNA localization in addition to translation regulation and mRNA turnover (Berleth et al., 1988; Clouse et al., 2008; Lin et al., 2006; Wilhelm et al., 2003).

Vap33 and dTtc1 (CG14894) were also enriched in the Egl interactome. We were interested in these proteins because their human orthologs, VAPB and TTC1, were known to interact (Lotz et al., 2008; Taipale et al., 2014). In addition, human TTC1 is present in the Dynein interactome (Redwine et al., 2017). Human and fly Ttc1 contains three tandem tetratricopeptide repeat (TPR) motifs, domains typically involved in scaffolding protein-protein interactions (Zeytuni and Zarivach, 2012). We therefore hypothesize that Ttc1 is a conserved cargo adaptor for Dynein, and is perhaps linked to Dynein via its interaction with the Egl/BicD complex.

The primary function attributed to Egl is as an RNA adaptor for Dynein (Dienstbier et al., 2009; Nashchekin and St Johnston, 2009). It was therefore somewhat surprising to find it in a complex with Vap33 and dTtc1. Human VAPB is an ER protein that serves to link membranes of the ER with other organelles (Dudas et al., 2021; Kamemura and Chihara, 2019). In addition, fly Vap33 is required for ER to Golgi trafficking (Mao et al., 2019). Our interactome analysis of dTtc1 revealed an enrichment of factors that localize to the endomembrane system. Why might Egl be interacting with proteins that are likely associated with vesicles? One possibility is that Egl has Dynein-independent functions and its interaction with Vap33 and dTtc1 have nothing to do with Dynein. Another possibility is that vesicles and mRNAs might be co-transported by the Egl/Dynein complex into the oocyte. This phenomenon of mRNA and vesicle co-transport has been recently observed in rat primary cortical neurons where mRNAs encoding stress response proteins co-traffic along with lysosomes (Liao et al., 2019). Consistent with a co-transport model, our results indicate that vesicles of the ER are highly enriched in the oocyte in an Egl-dependent manner (Fig. 6). During egg chamber maturation, oocyte volume expands greatly. Growth of the oocyte requires the almost unidirectional transport transport of mRNAs, proteins and vesicles from the nurse cells into the oocyte (Lu et al., 2021). An efficient mechanism of delivering these diverse cargo into the oocyte would be to co-transport particles containing multiple types of cargo, the biological equivalent of car pooling. Future studies will be aimed at testing this model of mRNA/vesicle co-transport.

Although we were able to recover BicD and Dlc in the Egl interactome, components of the Dynein motor were not enriched in the mass spectrometry data set (Fig. 3). By contrast, western blotting, a more sensitive approach, indicates that Egl-Trbo is capable of biotinylating Dhc (Fig. 2). One reason for this discrepancy might be that only a small percentage of Egl-Trbo is in a complex with Dynein. If this is true, Dynein components might not be heavily biotinylated and therefore not efficiently precipitated. Another possibility is that although Egl-Trbo abundantly associates with Dynein, the organization of the native complex precludes effective biotinylation of Dynein. BioID, the enzyme from which Trbo is derived, is thought to have a biotinylation radius of around 10 to 15nm (Kim et al., 2014). Dynein might be beyond this labeling radius to be effectively biotinylated by Egl-Trbo. A similar scenario was observed in mammalian cells. When the p62 component of the Dynactin complex was tagged with BioID, most components of the Dynactin complex were enriched in the p62-BioID pellet. However, components of the Dynein motor were not (Redwine et al., 2017). In addition, the interactome of BicD2 was significantly different depending on whether the tag was placed on the N or C terminus of the protein (Redwine et al., 2017). Given the high degree of co-localization observed between Egl and Dynein (Navarro et al., 2009), we favor the latter explanation; i.e. the limited labeling radius prevents effective biotinylation of Dynein by Egl-Trbo.

### Defining protein interactomes using nanobodies

Another objective of our study was to develop a nanobody-based approach for defining interactomes. We used the nanobody against GFP, referred to as the GFP binding protein (GBP), for this strategy. This nanobody was chosen because of its usefulness for the *Drosophila* community. Thousands of publicly available GFP tagged fly strains have been generated over the past several years (Buszczak et al., 2007; Kelso et al., 2004; Nagarkar-Jaiswal et al., 2015; Sarov et al., 2016). In addition, as we did with dTtc1, most labs wishing to examine the localization of a POI, often create a GFP tagged version of the gene. Thus, if GBP-Trbo is proven effective, any lab wishing to determine the interactome of their GFP-tagged protein would simply have to perform a single cross. Individual transgenes will not have to be generated for each POI.

To validate this approach, we determined the interactomes of endogenously tagged GFP-Me31b and UAS-driven GFP-dTtc1. Me31b is a component of P bodies and past studies have identified some of its interacting partners (DeHaan et al., 2017; Liu et al., 2011; McCambridge et al., 2020; Nakamura et al., 2004; Wilhelm et al., 2005). In total, 46 interactions have been reported for Me31b. GBP-Trbo was able to identify 21 of these interactions (Supplemental table 3). Our inability to detect the remaining interactions might have to do with the labeling radius caveat mentioned above. Another limitation relates to the availability of exposed lysine residues. If a protein that interacts with Me31b has few or no surface-exposed lysine residues, it will not be efficiently biotinylated regardless of its proximity to Me31b. These proteins will also be under-represented in the interactome. Regardless of these caveats, we were able to identify 293 unique interactions for Me31b. Consistent with the function of P bodies, GO analysis indicates a significant enrichment of proteins with known roles in post-transcriptional gene regulation (Fig. 8).

### Limitations and improvements of this strategy

In an ideal scenario, the expression of GBP-Trbo will be stoichiometric with the GFP tagged POI. In theory, this can be accomplished using a conditionally stabilized GBP (Tang et al., 2016). This version of GBP is targeted for proteosomal degradation if it is not bound to GFP, thus reducing background biotinylation. Our construct contained the destabilizing mutations. However, for reasons that we do not understand, GBP-Trbo was not degraded in the absence of the GFP tagged POI. Despite this issue, as long as an appropriate GFP tagged control is used, background biotinylation can be accounted for and removed from further analysis.

The current version of GBP-Trbo will bind to cytosolic POIs. However, if the GFP tag on a POI if present within the lumen of an organelle, it will not be accessible to GBP-Trbo. However, this limitation can be overcome by attaching an organelle targeting sequence to GBP-Trbo. Another potential issue is that cells expressing the GFP tagged POI might be present in a mix of cells, many of which are not GFP positive. In this case, FACS sorting might have to be used to isolate the GFP positive cells prior to protein purification.

As noted previously, a caveat of proximity biotinylation is the labeling radius of the enzyme. In theory, the labeling radius can be increased by lengthening the flexible linker between the POI and Trbo. The Roux lab demonstrated that a longer linker is indeed capable of identifying more distal components of a given complex (Kim et al., 2016). However, longer linkers might also compromise the function of the tagged protein. In this regard, GBP-Trbo might be more amenable to modification. Because the nanobody is a small globular protein, its binding activity is unlikely to be severely compromised by the addition of longer linkers. Care should be taken in using very long linkers, however, as this might also increase the rate of false-positive hits.

One way to increase the sensitivity of this approach would be to adapt the Moontag (or Suntag) approach to Trbo (Boersma et al., 2019). In this case, instead of fusing GBP to Trbo, the nanobody corresponding to Moontag (gp41 nanobody) is used. The recognition peptide for this nanobody is then fused to the POI in multiple copies, usually between 12 to 24 copies. When both constructs are expressed it the same cell, multiple copies of Trbo will be bound to the POI. This would greatly increase the level of biotinylation and subsequent enrichment of biotinylated proteins. This approach might facilitate the identification of proteins that are part of the interactome but not efficiently labeled by a single copy fo Trbo. Of course, a limitation of this approach is that it would require the generation of new transgenes for each POI.

## ACKNOWLEDGEMENTS

We would like to thank A. Nakamura for providing the Me31b antibody and H. Bellen for the Vap33 antibody. We are grateful to the Bloomington Stock Center, Developmental Studies Hybridoma Bank, and Addgene for providing fly strains, antibodies, cell lines, and DNA constructs. This work was supported by a grant from the National Institutes of Health to G.B.G (R01GM100088).

## MATERIALS AND METHODS

### DNA constructs and fly stocks

The Egl-Trbo and GFP-Trbo transgenes were constructed by cloning the cDNA for Egl or EGFP into the Kpn1 and BamH1 sites of the pUASp-attB-K10 vector (Koch et al., 2009). The Egl cDNA used contained point mutations that made it refractory to the *egl* shRNA (Sanghavi et al., 2016). The reverse translation of the published Trbo sequence (Branon et al., 2018) was gene synthesized using *Drosophila* codon optimization (Genewiz). The gene synthesized fragment also contained a 25 amino acid linker at the N-terminal end of the Trbo sequence as well as BamHI and XbaI restriction sites. This fragment was cloned downstream of Egl or EGFP. The sequence of this construct is listed in Supplemental Fig. 2. Both constructs were injected into Bloomington stock number 34760 (P{CaryIP}su(Hw)attP1, donor Nobert Perrimon) using PhiC mediated integration.

The GBP-Trbo construct was also cloned into pUASp-attB-K10. The above Trbo construct was cloned into the BamHI and XbaI sites of the vector. Two tandem copies of the GFP nanobody (GBP) separated by a 15 amino acid linker was generated by gene synthesis (Genewiz). The sequence also contained the mutations that were predicted to destabilize the nanobody in the absence of antigen binding (Tang et al., 2016). A 25 amino acid linker separated GBP and Trbo. The fragment also contained Kpn1 and BamH1 restriction sites. This gene synthesized product was cloned upstream of Trbo. The sequence of GBP-Tbro present within this construct is listed in Supplemental Fig. 3. This construct was injected into Bloomington stock number 24485 (M{3xP3-RFP.attP’} ZH-68E, donors Konrad Basler & Johannes Bischof).

The GFP-dTtc1 construct was also cloned into the pUASp-attB-K10 vector. The sequence for dTtc1 (CG14894) was reverse translated and generated by gene synthesis using *Drosophila* codon optimization (Genewiz). This construct also contained BamHI and XbaI restriction sites and was cloned into a vector containing EGFP. This construct was injected into Bloomington stock number 244749 (M{3xP3-RFP.attP} ZH-86Fb, donors Konrad Basler & Johannes Bischof). All fly strains were injected by BestGene Inc.

Endogenous Egl was depleted in the female germline using *egl* shRNA-1 (Bloomington stock center; #43550, donor TRiP). shRNA expression was driven using P{w[+mC]=matalpha4-GAL-VP16}V37 (Bloomington Stock Center, #7063; donor Andrea Brand) for early-stage expression. GFP-tagged Golgi vesicles were visualized by crossing w[*]; P{w[+mC]=UASp-GFP.Golgi}1 (Bloomington Stock Center, #30902; donor Richa Rikhy) with the above Gal4 driver. Fly crosses used for all experiments were maintained at 25°C.

The GFP-Snx9 construct was generated by cloning the cDNA for Snx9 (Hicks et al., 2015) into the HindIII and EcoR1 sites of pEGFP-C3 (Clontech). The mammalian Trbo-GBP construct was generated by cloning a gene synthesize fragment containing two tandem copies of the GFP nanobody into the V5-TurboID-NES_pCDNA3 vector. V5-TurboID-NES_pCDNA3 was a gift from Alice Ting (Addgene plasmid # 107169 ; http://n2t.net/addgene:107169 ; RRID:Addgene_107169) (Branon et al., 2018). The insert was cloned using Gibson assembly (NEB) immediately downstream of Trbo. The sequence of this construct is listed in Supplemental Fig. 4.

Fly strains used in this study will be deposited at the Bloomington *Drosophila* Stock Center. DNA constructs used in this study will be deposited at Addgene.

### Protein purification

Flies were grown on standard media (Per 1L, 83.3ml molasses, 6.7g agar, 9g dry yeast, 68.3g cornmeal and 4.18ml propionic acid) without adding exogenous biotin. In our experience, this media formulation contains adequate levels of biotin such that the expression of Trbo enables the efficient biotinylation of proteins as determined using Streptavidin conjugated Alexa647 (for localization studies) or Streptavidin conjugated HRP (for western blotting). Small scale binding experiments were performed using 1mg of ovarian lysate. Lysates were prepared by homogenizing ovaries in RIPA buffer (50 mM Tris-Cl [pH 7.5], 150 mM NaCl, 1% NP-40, 1 mM EDTA) containing a Halt Protease inhibitor cocktail (Piece). Biotionylated proteins were precipitated by incubating in 15ul of High Capacity Streptavidin Agarose beads (Piece). Binding was performed overnight at 4°C. The next day the sample were washed 4 times with 1ml RIPA. Bound proteins were eluted in Laemmli buffer, run on a gel, and analyzed by western blotting. All western blot images were acquired on a Bio Rad ChemiDoc MP.

For proteomics experiments, 85 ovaries were dissected for each genotype. Approximately 5.5mg of ovarian lysates were used in each binding experiment. Lysates were prepared as described above. Biotionylated proteins were purified by incubating in 50ul of High Capacity Streptavidin Agarose beads at 4°C overnight. The next day the samples were extensive washed using 1ml of the following solutions: 3 times with RIPA buffer, 3 times with 1% SDS, 3 times with RIPA buffer, 3 times with high salt RIPA buffer (50 mM Tris-Cl [pH 7.5], 1M NaCl, 1% NP-40, 1 mM EDTA), 2 times with RIPA buffer and 4 times with PBS. The final washes in PBS are required to remove as much of the detergent as possible as this will affect the downstream mass spectrometry analysis. The entire experiment was done in triplicate for proteomics analysis.

### Mass spectrometry

The mass spectrometry to determine the Egl interactome (Fig. 3) was performed at the Augusta University proteomics core. The following protocol was used. The beads with bound proteins were reduced with dithiothreitol, alkylated using iodoacetamide in 8M urea denaturation buffer (50mM Tris-HCl, pH 8) and digested overnight in 50mM ammonium bicarbonate using trypsin (Thermo Scientific #90057) at 37°C. Digested peptides were cleaned using a C18 spin column (Harvard Apparatus #744101) and then lyophilized. Peptide digests were analyzed on an Orbitrap Fusion tribrid mass spectrometer (Thermo Scientific) coupled with an Ultimate 3000 nano-UPLC system (Thermo Scientific). Two microliters of reconstituted peptide was first trapped and washed on a Pepmap100 C18 trap (5um, 0.3×5mm) at 20ul/min using 2% acetonitrile in water (with 0.1% formic acid) for 10 minutes and then separated on a Pepman 100 RSLC C18 column (2.0 um, 75-μm × 150-mm) using a gradient of 2 to 40% acetonitrile with 0.1% formic acid over 40 min at a flow rate of 300nl/min and a column temperature of 40°C.

Samples were analyzed by data-dependent acquisition in positive mode using Orbitrap MS analyzer for precursor scan at 120,000 FWHM from 300 to 1500 m/z and ion-trap MS analyzer for MS/MS scans at top speed mode (3-second cycle time). Collision-induced dissociation (CID) was used as fragmentation method. Raw data were processed using Proteome Discoverer (v1.4, Thermo Scientific) and submitted for SequestHT search against the Uniprot *Drosophila* database. Percolator PSM validator algorithm was used for peptide spectrum matching validation. SequestHT search parameters were 10 ppm precursor and 0.6 Da product ion tolerance, with static Carbamidomethylation (+57.021 Da).

The GBP-based proteomics experiments (Figs. 8 and 9) were performed by the Emory Integrated Proteomics Core. This core was used because the mass spectrometer at our institution was being serviced at the time and was unavailable for use. For on-bead digestion, a published protocol was followed (Soucek et al., 2016). To the beads, digestion buffer (50 mM NH4HCO3) was added and the mixture was then treated with 1 mM dithiothreitol (DTT) at room temperate for 30 minutes, followed by 5 mM iodoacetimide (IAA) at room temperate for 30 minutes in the dark. Proteins were digested with 0.5ug of lysyl endopeptidase (Wako) at room temperate overnight and were further digested overnight with 1ug trypsin (Promega) at room temperate. Resulting peptides were desalted with HLB column (Waters) and were dried under vacuum.

The data acquisition by LC-MS/MS was adapted from a published protocol (Seyfried et al., 2017). Derived peptides were resuspended in the loading buffer (0.1% trifluoroacetic acid, TFA) and were separated on a Water’s Charged Surface Hybrid (CSH) column (150um internal diameter (ID) x 15 cm; particle size: 1.7um). The samples were run on an EVOSEP liquid chromatography system using the 30 samples per day preset gradient and were monitored on a Q-Exactive Plus Hybrid Quadrupole-Orbitrap Mass Spectrometer (ThermoFisher Scientific). The mass spectrometer cycle was programmed to collect one full MS scan followed by 20 data dependent MS/MS scans. The MS scans (400-1600 m/z range, 3 × 106 AGC target, 100 ms maximum ion time) were collected at a resolution of 70,000 at m/z 200 in profile mode. The HCD MS/MS spectra (1.6 m/z isolation width, 28% collision energy, 1 × 105 AGC target, 100 ms maximum ion time) were acquired at a resolution of 17,500 at m/z 200. Dynamic exclusion was set to exclude previously sequenced precursor ions for 30 seconds. Precursor ions with +1, and +7, +8 or higher charge states were excluded from sequencing.

Label-free quantification analysis was adapted from a published procedure (Seyfried et al., 2017). Spectra were searched using the search engine Andromeda, integrated into MaxQuant, against *Drosophila melanogaster* (Fruit fly) Uniprot database (42,676 target sequences). Methionine oxidation (+15.9949 Da), asparagine and glutamine deamidation (+0.9840 Da), and protein N-terminal acetylation (+42.0106 Da) were variable modifications (up to 5 allowed per peptide); cysteine was assigned as a fixed carbamidomethyl modification (+57.0215 Da). Only fully tryptic peptides were considered with up to 2 missed cleavages in the database search. A precursor mass tolerance of ±20 ppm was applied prior to mass accuracy calibration and ±4.5 ppm after internal MaxQuant calibration. Other search settings included a maximum peptide mass of 6,000 Da, a minimum peptide length of 6 residues, 0.05 Da tolerance for orbitrap and 0.6 Da tolerance for ion trap MS/MS scans. The false discovery rate (FDR) for peptide spectral matches, proteins, and site decoy fraction were all set to 1 percent. Quantification settings were as follows: re-quantify with a second peak finding attempt after protein identification has completed; match MS1 peaks between runs; a 0.7 min retention time match window was used after an alignment function was found with a 20-minute RT search space. Quantitation of proteins was performed using summed peptide intensities given by MaxQuant. The quantitation method only considered razor plus unique peptides for protein level quantitation.

### Antibodies

The following antibodies were used: anti-FLAG (Sigma Aldrich, F1804, 1:5000 for western, 1:500 for immunofluorescence), mouse anti-BicD (Developmental Studies Hybridoma Bank, clones 1B11 and 4C2, 1:300 for western, donor R. Steward), rabbit anti-Ctp (Abcam, ab51603, 1:5000 for western), mouse anti-GFP (Clontech, JL-8, 1:5000 for western), mouse anti-Dhc (Developmental Studies Hybridoma Bank, clone 2C11-2, 1:300; for western, donor J. Scholey), rabbit anti-Me31b (Gift of A. Nakamura, 1:12000 for western, 1:1200 for immunofluorescence), rabbit ant-Vap33 (Gift of H. Bellen, 1:10000 for western, 1:2000 for immunofluorescence), mouse anti-Cnx99A (Developmental Studies Hybridoma Bank, clone Cnx99A 6-2-1, 1:60 for immunofluorescence, donor S. Munro), mouse anti-Golgin-84 (Developmental Studies Hybridoma Bank, clone Golgin84 12-1, 1:60 for immunofluorescence, donor S. Munro) (Riedel et al., 2016), rabbit anti-RFP (1:2000 for immunofluorescence)(Goldman et al., 2019), mouse anti-BicD2 (Thermo Fisher, clone MA5-23522, 1:1000 for western) and mouse anti-V5 (Thermo Fisher, catalog R960-25, 1:1000 for immunofluorescence). The following secondary antibodies were used: goat anti-rabbit Alexa 594, 555 and 488 (Life Technologies, 1:400, 1:400 and 1:200 respectively); goat anti-mouse Alexa 594, 555 and 488 (Life Technologies, 1:400, 1:400 and 1: 200 respectively) goat anti-mouse HRP (Pierce, 1:5000); and goat anti-rabbit HRP (Pierce, 1:5000).

### Immunofluorescence

Immunofluorescence was performed as previously described (Goldman et al., 2019). In brief, ovaries were dissected and fixed in 4% formaldehyde (Pierce) for 20 min at room temperature. The primary antibody was incubated in blocking buffer (PBS + 0.1% Triton X-100 + 2% BSA) overnight at 4°C. Next, the samples were washed three times in PBST (PBS + 0.1% Triton X-100) and incubated overnight with the fluorescent secondary antibody in the same blocking buffer. The samples were then washed four times with PBST, stained with DAPI, and mounted onto slides using Prolong Diamond (Life technologies). For detecting biotinylated proteins, the samples were incubated with Streptavidin Alexa647 (Life technologies; 1:1200). The Strep674 was added to the sample at the same time as the secondary antibody.

### Microscopy

Images were captured on either a Zeiss LSM 780 inverted confocal microscope or an inverted Leica Stellaris confocal microscope. Images were processed for presentation using Fiji, Adobe Photoshop, and Adobe Illustrator. All imaging experiments were performed at the Augusta University Cell Imaging Core.

### Quantification

Protein enrichment within the oocyte was quantified by measuring the average pixel intensity of the localized signal and dividing by the average pixel intensity of the delocalized signal. ImageJ/Fiji was used for this analysis. Graphs were assembled using Graphpad Prism9.

### Illustrations

The egg chamber illustration in Fig. 1 was created with BioRender.com. The illustration in Fig.2 was made using Adobe Illustrator.

## FIGURE LEGENDS

**Supplemental Figure 1: (A)** Egg chambers expressing a control shRNA against *eb1* (A) or an shRNA against *egl* (A’) were fixed and processed for immunofluorescence using an antibody against Me31b. Me31b signal is shown using a red to white LUT. Although P bodies are present in Egl depleted egg chambers, diffuse cytoplasmic Me31b localization is increased. **(B)** HeLa cells were transiently transfected with constructs expressing either Trbo alone or Trbo-TTC1. Lysates were prepared from these cells and biotinylated proteins were purified using streptavidin agarose. Bound proteins were eluted and analyzed by western blotting using an antibody against BicD2. **(C)** Egg chambers were fixed and processed for immunofluorescence using antibodies against Vap33 (red) and Cnx99A (green), a marker for ER vesicles. This image show the germarium localization of Vap33 and ER. The scale bar is 20 microns.

**Supplemental Figure 2:** Sequence of Trbo used in these studies.

**Supplemental Figure 3:** Sequence of GBP-Trbo used for expression in *Drosophila* tissue.

**Supplemental Figure 4:** Sequence of Trbo-GBP used for expression in mammalian cells.

**Supplemental Table 1:** List of proteins that interact with Egl.

**Supplemental Table 2:** List of proteins that interact with Me31b.

**Supplemental Table 3:** List of proteins that were known to interact with Me31b from previous studies. This list also indicates which of these proteins were recovered in our study.

**Supplemental Table 4:** List of proteins that interact with dTtc1.

